# Beyond gene length: Exon-intron architecture and isoform potential in the evolution of eukaryotic complexity

**DOI:** 10.64898/2026.03.25.714307

**Authors:** Senbao Lu, Yue Bao, Gloria M. Sheynkman, Dmitry Korkin

## Abstract

Alternative splicing is a major source of human transcriptomic and phenotypic variation, yet its evolutionary contribution to genomic complexity remains unresolved. It has been shown that a mean gene length can be the most basic but remarkably efficient proxy for multicellular genome complexity, whereas mean protein length is not, as it plateaus abruptly early in eukaryotic evolution. Here, we show that, across 2,683 genomes, exon count continues to increase beyond this transition and then rapidly saturates at ∼10 exons per gene, supporting its role as an additional dimension of genomic complexity linked to exon-intron architecture. A simple stochastic exon-splitting model reproduces the observed biphasic exon-number growth pattern and identifies minimal exon length as a key determinant.

## Background

Both coding and non-coding regions of a gene contribute to organismal complexity throughout the course of eukaryotic evolution [1]. Yet how their relative contributions change over evolutionary time, and whether increasing complexity is driven primarily by regulatory architecture or by coding potential is still an open question. The emergence of introns in eukaryotes created an opportunity for controlled alternative splicing, leading to a major expansion of complexity in multicellular organisms [1, 2]. Although the exon-intron architecture of eukaryotic genes is widely recognized as a major contributor to genomic complexity of the eukaryotic genomes, the extent to which alternative splicing contributes to proteomic variation and, ultimately, phenotypic diversity remains debated [3].

A recent work by Muro *et al* [4] studied the distribution of protein and gene lengths across over 6,500 species, showing that mean protein length grows following a two-phase pattern. In the first phase, mean protein length increases with mean gene length, while in the second phase, which begins at a mean gene length of approximately 1,500 bp, mean protein length abruptly plateaus at around 500 amino acid residues and no longer scales with gene length. Subsequent increases in gene length are therefore largely attributable to non-coding components of gene architecture, including introns and untranslated regions (UTRs). With the length of coding sequence no longer being a proxy for organismal complexity does it imply that the complexity of multicellular organism later in the eukaryotic evolution is driven exclusively by the noncoding part of the gene? The emergence of alternative splicing argues otherwise. A gene with a coding sequence of 1,500 bp distributed across ten exons has a substantially greater capacity to generate multiple protein-coding isoforms than a gene of the same coding length composed of a single exon, thereby breaking the ‘one gene-one protein’ paradigm and expanding functional diversity of the proteome.

In this Brief Report, we complement the study of Muro *et al*. [4] by analyzing how the average exon count per genome scales with average gene length across the full spectrum of eukaryotic genomes and by examining the relationship between protein-coding isoform number and exon count in six model genomes. We further propose a simple stochastic model of exon-number growth per gene, which suggests that this process is fundamentally constrained by minimal exon length.

## Results and Discussion

First, we examined the relationship between the mean number of exons per gene and its standard deviation using a comprehensive set of 2,683 genomes extracted from Ensembl (release 114) [5]. We considered genomes with mean exon count for the canonical isoform greater than 1.0, indicating the presence of alternative splicing. The earliest such genome (*Kipferlia bialata*) had a mean gene length of 681 bp, however the average exon count began to rise sharply at the mean gene length of ∼1,500 bp. We found the relationship between the mean exon count and its standard deviation was linear, *S* = 1.09*m* − 1, (Fig. 1A) and followed the mean-variance scaling relationship, described by Taylor’s law, a general statistical law between the mean *m*, and standard deviation, *s*, (or variance, *v*) frequently seen in complex biological systems: *v* = *s*^*2*^ = *am*^*b*^ [6]. The nearly perfect fit of this relationship (R^2^ = 0.951) implied a nearly universal coefficient of variation of exon numbers across eukaryotes. Furthermore, the exponent *b* ≈ 2 observed in our data had been previously observed across various proteomic and transcriptomic datasets, including protein concentration fluctuations [7], single-cell transcriptomic expression [8], and bulk RNA-Seq counts [9]. The exon-number variability nearly linear growth with the mean was consistent with a multiplicative evolutionary process shaping intron-exon architecture across eukaryotic genomes.

**Figure 1.**
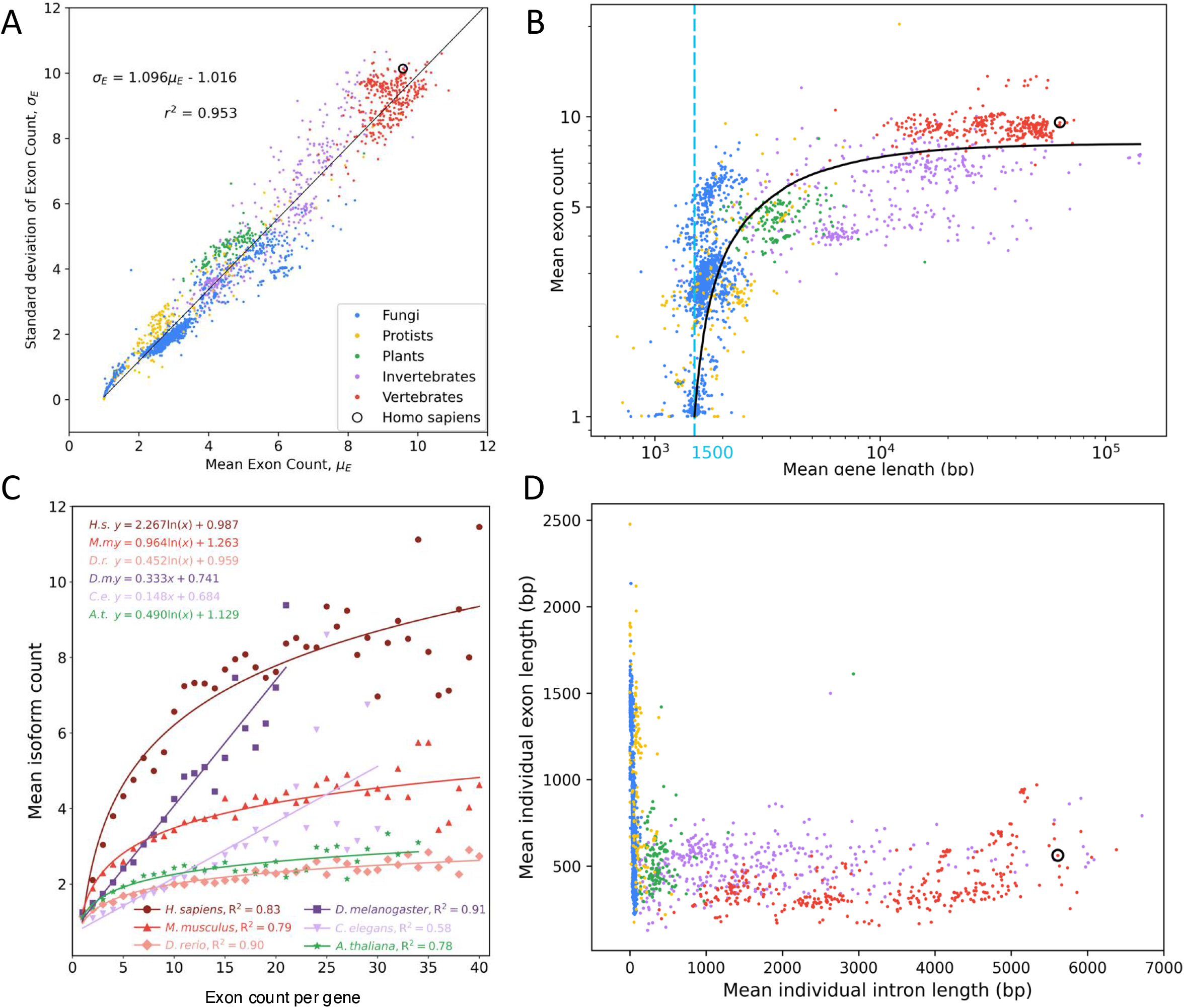
Evolution of genomic complexity characterized by a mean exon count. **A**: Relationship between the mean exon count per gene and its standard deviation across 2,683 eukaryotic genomes, showing a near-perfect linear fit consistent with Taylor’s law. Black line indicates the optimal linear fit. A data point corresponding to human is circled. **B**: Mean exon count per gene continues to increase after protein length reaches a plateau at ∼1,500 bp, revealing a biphasic pattern: rapid growth in fungi and early protists followed by markedly slower growth in plants, invertebrates, and vertebrates, with saturation at ∼10 exons per gene. The black line shows the scaling behavior predicted by the stochastic exon-splitting model. A data point corresponding to human is circled. **C**: Best-fitting relationships between isoform count and exon count based on comparisons of logarithmic and linear models. Plant and vertebrate model organisms were better described by a logarithmic fit. **D**: Exon and intron length distributions differ markedly between fungi and protists (blue and yellow) and plants, invertebrates and vertebrates (green, purple and red), indicating distinct modes of exon–intron architectural expansion across eukaryotes. A data point corresponding to human is circled.

Second, we examined how the average exon count per gene scales with overall gene length. We found that, after mean protein length stabilizes at ∼1,500 bp [4], exon count continues to increase, undergoing a similar transition between two distinct phases: a rapid rise in fungi and early protists, followed by a markedly slower increase in plants, invertebrates, and, ultimately, vertebrates (Fig. 1B). Because exon number can in theory expand the space of potential isoforms exponentially, we hypothesized that the apparent plateau in exon-intron architectural complexity at approximately 10 exons per gene, on average, may reflect the attainment of sufficient isoform-generating capacity to support the complex functions of higher vertebrates. Consistent with this idea, when we analyzed the average number of isoforms per gene as a function of exon count across the transcriptomes of six model organisms and compared logarithmic and linear regression fits (Fig. 1C, Supplementary Table 1C), four species, including human, mouse, zebrafish, and Arabidopsis, showed a substantially better logarithmic fit (R^2^ = 0.78-0.90), whereas the two invertebrates, fruit fly and nematode, were slightly better described by a linear fit. Although this analysis was limited to species with the most comprehensively annotated isoform repertoires, the results suggest that, in vertebrates and plants, isoform number increases significantly slower than its theoretical combinatorial potential. Notably, based on the scaling coefficients of the logarithmic fits (Fig. 1C), the human transcriptome appears to use the largest number of isoforms per gene on average, thereby maximizing the potential contribution of alternative splicing to proteomic and, consequently, functional diversity.

Analysis of the length distributions of individual exons and introns showed that, consistent with previous observations, their growth dynamics differ markedly between fungi and protists, on the one hand, and plants, invertebrates, and vertebrates, on the other (Fig. 1D). In the former group of genomes, the expansion of gene architecture occurs primarily through elongation of individual exons in the presence of short introns, whereas in the latter group, the pattern is reversed, with substantial expansion of individual introns while exon length remains constrained. Interestingly, the growth of individual introns within exon-intron gene architecture continues despite the increasing number of exons and introns per gene, suggesting that multicellular complexity is shaped not only by overall expansion of the non-coding portion of the gene, but also by the increase in average intron length.

Finally, we found that the scaling of exon count with gene length can be recapitulated by a simple stochastic exon-splitting model defined by two parameters: the rate of exon-splitting events and the minimal exon length. By optimizing these parameters to achieve the best fit to the available data (Fig. 1B), we estimated the minimal exon length to be ∼ 138-139 bp, corresponding to a protein length of ∼46 amino acid residues, somewhat larger than the previous estimate of ∼50 bp [10]. Notably, our value is comparable to the proposed minimal size of a protein domain, ∼40-50 residues [11, 12]. Together, these observations suggest two, not mutually exclusive, factors that may contribute to the limited exon growth in the higher eukaryotes: (1) the attainment of sufficient isoform-generating capacity, and (2) the requirement for a minimal exon length in the context of an already stabilized overall protein length.

## Conclusions

With the unprecedented expansion of large-scale omics data over the past two decades, recurring patterns have emerged that can be described by common mathematical laws. Log-normal and power-law distributions in biological networks [13-15] and Taylor’s law [6], connecting mean and standard deviation of the data first in ecology, and later in proteomics and transcriptomics [7-9] are well-studied examples of such patterns. Our work, which quantifies the basic exon-intron architecture complexity across eukaryotes, also finds previously observed patterns. First, the observed linear relationship between the standard deviation and the mean exon count implies a quadratic scaling of variance with the mean, consistent with Taylor’s law. Likewise, the relationship between mean gene length, another basic measure of genomic complexity, and its variance, which emerges from multiplicative processes of gene growth, also follows Taylor’s law [4]. In this context, our findings may suggest that as genes acquire more exons on average, exon-number diversification accelerates disproportionately, consistent with a non-additive expansion of gene architectural complexity.

Second, similar patterns emerge when comparing the scaling of protein length and exon count as functions of average gene length. In both cases, the relationship can be described by two phases: an initial phase of rapid growth followed by a plateau. Notably, however, the rapid growth phase of exon count continues even after protein length has entered its plateau phase. Together, these results suggest that exon-intron architecture captures an additional dimension of genomic complexity that is not reflected by the lengths of gene’s coding and non-coding regions alone, adding to the growing body of evidence that introns and exons make coupled contributions to the complexity and diversity of multicellular genomes [1, 16, 17]. Furthermore, they support a view of eukaryotic complexity, in which continued expansion of gene architecture, and the isoform-generating potential it enables, remains a major determinant of multicellular complexity alongside the expansion of the non-coding regions, even after coding length itself has stabilized.

## Methods

In this study, we aim to analyze how an average exon count and isoform count per gene change as functions of average gene length across the full spectrum of eukaryotic genomes. To achieve this goal, we first compile gene architecture information for each genome and extract the corresponding numbers characterizing genes, their exons, and isoforms. Then, we perform statistical analysis to understand how exon count changes across different phyletic groups. Finally, we propose a stochastic model of exon splitting to explain the exon growth pattern.

### Gene Annotations / Data collection

We collect 2,683 genomes for species in fungi, protists, plants, invertebrates, and vertebrates from EnsemblGenome (release 61) and Ensembl (release 114) [5]. Specifically, we download the GFF3 (General Feature Format) files of 1,501 fungi, 240 protists, 225 plants, and 374 invertebrates from EnsemblGenome FTP site, and 343 vertebrates from Ensembl FTP site. For each species, we extract gene length and canonical transcript ID for each gene, as defined by Ensembl’s ‘Canonical’ tag. Since there is only one canonical transcript per gene, we further extract exon count and length, CDS (coding sequence) count and length, and the lengths of 5’ UTR and 3’ UTR regions. We assign these values to the corresponding gene.

### Isoform Annotations for model organisms

For the six model organisms, namely *Arabidopsis thaliana* of plant, *Caenorhabditis elegans* and *Drosophila melanogaster* of invertebrates, and *Danio rerio, Mus musculus*, and *Homo sapiens* of vertebrates, we additionally extract the total isoform count from the GFF3 files. These organisms are selected because their transcriptomes have been extensively characterized, providing complete or near-complete isoform annotations in Ensembl, whereas most other Ensembl genomes contain incomplete isoform information. For comparative analysis and curve fitting, genes from each of the six genomes are aggregated based on the number of exons they have into data points (*x*_*i*_, *y*_*i*_), where *x*_*i*_ corresponds to the exon count, and *y*_*i*_ corresponds to the mean isoform count over all genes containing *x*_*i*_ exons. To enable comparison across organisms, we restricted the analysis to genes with exon counts between 1 and 40. In addition, data points for which fewer than 10 genes were available at a given *x*_*i*_ were excluded from the analysis. We also excluded a single outlier data point in *Drosophila melanogaster* corresponding to *x* = 24 exons and a mean isoform count of *y* ≈ 15. We then fit two functions independently for each species, linear and logarithmic, and select the model with the higher R^2^ coefficient value as the final fit.

### The stochastic model of exon count growth

Last, we introduce a stochastic model to explain a two-phase like behavior of the exon number growth with respect to average gene length, characterized by rapid exon count growth initially followed by plateauing of this number. The model is based on two basic assumptions that (i) exon splits into two exons at a constant rate, and (ii) there exists a minimum exon length for efficient splicing. While the second assumption has been independently proposed and evaluated [10, 18], the first assumption is made due to our limited knowledge on the rate of evolutionary events that increase the number of exons, such as splicing mutations [19]. Based on these assumptions, the stochastic model has two parameters: the minimum exon length *l*_*min*_, and a probability of exon splitting, ε.

The model consists of two basic steps:

#### Step 1 (Initialization)

We initialize the stochastic process using *N* genes, each gene containing 1 CDS of length 1,500 nt. Each CDS initially contains 1 exon of length 1,500 nt. We not that in this model UTR regions are not considered, as our analysis indicates that their average lengths, for both 3’ and 5’ UTRs plateau very quickly, compared to the exon length (see Suppl. Fig S1). We record the exon lengths for each gene as a vector of integers. The initial exon length list is thus *L*_*i*_ (0) = [1,500], *i*= 1, 2, …, *N*.

#### Step 2 (Iteration)

At each iteration step of the algorithm, denoted as *s* = 1, 2,…, one gene *k* is randomly selected, with its current vector of exon lengths being *L*_*k*_ (*s*−1). A random number ζ ∈[0,1] is then generated from a uniformly distributed probability density function *f*(ζ). If ζ < ε, then we randomly select one exon from the list *L*_*k*_ (*s*−1), and randomly split that exon into two by selecting a random split position anywhere in that exon. If both newly generated exons are longer than the minimum exon length *l*_*min*_, we keep this split and update the exon length list as *L*_*k*_(*s*). Otherwise, the split is rejected and *L*_*k*_ (*s*) = *L*_*k*_ (*s*−1). The exon length lists of the remaining *N*−1 genes remain unchanged: *L*_*i*_ (*s*) = *L*_*i*_ (*s*-1), ∀*i* ≠ *k*.

We run the simulation of this model with *N* = 2,000 genes for at most 100,000 iterations. The stoppage criterion for the simulation is defined based on the maximum average gene length in the collected data. Both parameters, *ε* and *l*_*min*_, are optimized through a grid search by minimizing the RMSE of the resulting fit of the generated exon count *vs*. gene length function. For the grid search the following protocol is used: for *l*_*min*_, we first run a grid search from 20 to 200 with an increment of 20 to find the optimized range, then search from 120 to 160 with an increment of 1 to find an integer that minimizes the RMSE. *ε* is related to step size defined in **Step 2**, as it represents the probability of exon splitting in evolutionary time. At the step size of 0.1 million years, *ε* is optimized at 0.001.

## Declarations

### Ethics approval and consent to participate

Not applicable

### Consent for publication

Not applicable

### Availability of data and materials

The datasets generated and/or analyzed during the current study are available in the GitHub repository, https://github.com/KorkinLab/AS_evolution.

### Author contributions

D.K. designed research; S.L. and Y.B. performed research; D.K., S.L., Y.B., and G.M.S. analyzed data and wrote the paper.

## Supplementary Materials

**Table S1.** Summary of logarithmic vs. linear curve fitting for six model organisms. The fit with the highest R^2^ score is shown in bold.

| Model organism | Linear function | Linear $R^2$ | Log function | Log $R^2$ |
| --- | --- | --- | --- | --- |
| *H. sapiens* | $y = 0.15x + 4.21$ | 0.63 | $y = 2.27\ln(x) + 0.99$ | **0.83** |
| *M. musculus* | $y = 0.06x + 2.64$ | 0.60 | $y = 0.96\ln(x) + 1.26$ | **0.79** |
| *D. rerio* | $y = 0.03x + 1.54$ | 0.82 | $y = 0.45\ln(x) + 0.96$ | **0.90** |
| *D. melanogaster* | $y = 0.33x + 0.74$ | **0.91** | $y = 2.32\ln(x) - 0.61$ | 0.77 |
| *C. elegans* | $y = 0.15x + 0.68$ | **0.58** | $y = 1.39\ln(x) - 0.49$ | 0.48 |
| *A. thaliana* | $y = 0.04x + 1.70$ | 0.67 | $y = 0.49\ln(x) + 1.13$ | **0.78** |

