## Supplementary Table S1 for "Beyond gene length: Exon-intron architecture and isoform potential in the evolution of eukaryotic complexity"

### Supplementary Materials

**Table S1. Summary of logarithmic vs. linear curve fitting for six model organisms.** The fit with the highest R^2^ score is shown in bold.

| **Model organism** | **Linear function** | **Linear R^2^** | **Log function** | **Log R^2^** |
| --- | --- | --- | --- | --- |
| *H. sapiens* | *y*= 0.15*x*+4.21 | 0.63 | y=2.27ln(x)+0.99 | **0.83** |
| *M. musculus* | *y*=0.06*x*+2.64 | 0.60 | y=0.96ln(x)+1.26 | **0.79** |
| *D. rerio* | *y*=0.03*x*+1.54 | 0.82 | y=0.45ln(x)+0.96 | **0.90** |
| *D. melanogaster* | *y*=0.33*x*+0.74 | **0.91** | y= 2.32ln(x)-0.61 | 0.77 |
| *C. elegans* | *y*=0.15*x*+0.68 | **0.58** | y= 1.39ln(x)-0.49 | 0.48 |
| *A. thaliana* | *y*=0.04*x*+1.70 | 0.67 | y= 0.49ln(x)+1.13 | **0.78** |
